# Query to reference single-cell integration with transfer learning

**DOI:** 10.1101/2020.07.16.205997

**Authors:** Mohammad Lotfollahi, Mohsen Naghipourfar, Malte D. Luecken, Matin Khajavi, Maren Büttner, Ziga Avsec, Alexander V. Misharin, Fabian J. Theis

## Abstract

Large single-cell atlases are now routinely generated with the aim of serving as reference to analyse future smaller-scale studies. Yet, learning from reference data is complicated by batch effects between datasets, limited availability of computational resources, and sharing restrictions on raw data. Leveraging advances in machine learning, we propose a deep learning strategy to map query datasets on top of a reference called *single-cell architectural surgery* (scArches, https://github.com/theislab/scarches). It uses transfer learning and parameter optimization to enable efficient, decentralized, iterative reference building, and the contextualization of new datasets with existing references without sharing raw data. Using examples from mouse brain, pancreas, and whole organism atlases, we showcase that scArches preserves nuanced biological state information while removing batch effects in the data, despite using four orders of magnitude fewer parameters compared to *de novo* integration. To demonstrate mapping disease variation, we show that scArches preserves detailed COVID-19 disease variation upon reference mapping, enabling discovery of new cell identities that are unseen during training. We envision our method to facilitate collaborative projects by enabling the iterative construction, updating, sharing, and efficient use of reference atlases.

## Introduction

Large single-cell reference atlases [1–4] comprising millions [5] of cells across tissues, organs, developmental stages, and conditions are now routinely generated by consortia such as the Human Cell Atlas [6]. These references help to understand the cellular heterogeneity that make up natural and inter-individual variation, ageing, environmental influences, and disease. We call such atlases “references” as their central goal is to enable users to learn from these comprehensive maps and thus analyse their own data (e.g., compare disease data to a healthy reference). Indeed, reference atlases provide an opportunity to radically change how we analyze single-cell datasets currently [6]: by learning from the appropriate reference we could automate the tedious clustering and annotation of new dataset, and easily perform comparative analyses across tissues, species, and disease conditions.

Learning from a reference atlas requires mapping a query dataset to this reference to generate a joint embedding. Yet, query datasets and reference atlases typically comprise data generated in different labs, with differing experimental protocols and thus contain batch effects. Data integration methods are typically used to overcome these batch effects in reference construction [7]. However, these approaches require access to all datasets used to generate the reference, which can be prohibitive especially for human data due to legal restrictions on data sharing. Furthermore, contextualizing a single dataset in this manner would require rerunning the full integration pipeline, which presupposes computational expertise and sufficient computational resources. Finally, traditional data integration methods assume that any perturbation between datasets that affects most cells is a technical batch effect to be removed. However, biological perturbations, such as disease, may also affect most cells. Thus, even if all above requirements are met, conventional approaches would not suffice for mapping query data onto references across biological conditions.

Exploiting large reference datasets is a well-established approach in Computer Vision [8] and Natural Language Processing [9]. In these fields, the commonly used deep learning approaches typically require a large number of training samples, which are not always available. By leveraging weights learned from large reference datasets to enhance learning on a target or query dataset [10], transfer learning (TL) models such as image-net [11] and BERT [12] have revolutionized analysis approaches [8, 9]: TL has improved method performance with small datasets (e.g., clustering [13], classification/annotation [14]) and enabled sharing of models using model zoos [15–17]. Recently, transfer learning has been applied on scRNA-seq for denoising [18], variance decomposition [19], and cell type classification [20, 21]. However, the current TL approaches in genomics show several drawbacks: they do not account for technical effects within and between reference and query[18], and lack systematic retraining with query data [19–21]. These limitations can lead to spurious predictions on query data with no, or small, overlap in cell types, tissues, species, or cell states [22, 23]. Given the recent success of deep learning models for data integration in single-cell genomics [7, 24–26], TL may also prove transformative to address the issues of efficient learning from reference data and model sharing.

We propose a novel TL and fine-tuning strategy for conditional neural network models called *architecture surgery*, implemented in the scArches pipeline. scArches is a fast and scalable tool for updating, sharing, and using reference atlases. Specifically, given a basic reference atlas, scArches enables users to share this reference as a trained network with other users, who can in turn update the reference using query-to-reference mapping and partial weight optimization without any sharing of their potentially private or unpublished data. Thus, users can build their own extended reference models, or perform stepwise analysis of datasets as they are collected, which is often crucial for emerging clinical datasets. Furthermore, scArches allows users to learn from reference data by contextualizing new, e.g. disease, data with a healthy reference in a joint latent space representation. Within this representation, one can perform reference-based clustering and transfer of annotation to query cells. We demonstrate the above features of scArches using publicly available scRNA-seq datasets ranging from pancreas to whole mouse atlases and COVID-19 immune cells. scArches is able to iteratively update a pancreas reference, transfer knowledge between mouse atlases, and map COVID-19 data onto a healthy reference while preserving disease-specific variation.

## Results

### Architectural surgery enables decentralized data integration by mapping novel query studies to reference datasets

Architectural surgery is a TL approach that adapts existing deep learning models. After training on multiple reference datasets, trained weights can subsequently be transferred with only minor weight adaptation (fine tuning) to map or overlay a novel study onto this reference. While this approach can be applied to any deep learning model, here we apply scArches to our recently proposed transfer variational autoencoder model trVAE [27] and to Negative Binomial conditional VAEs (CVAE) (see **Methods**) similar to scVI [25].

Consider the scenario with *N* ‘reference’ scRNA-seq datasets of a particular tissue or organism. We assign a categorical label *S*_*i*_ to each dataset that corresponds to the study label, which is encoded as a condition in the CVAE. These study labels may index traditional batch IDs (i.e., samples, experiments across labs, or sequencing technologies), biological batches (i.e., organs or species when used over the set of orthologous genes), perturbations such as disease, or a combination of these categorical variables.

We pretrain the model with the reference studies *S*_1:*N*_ (**Figure 1a**), which results in a latent space free of technical variation across these datasets [28]. Thus, we can use this embedding for further downstream analysis such as visualizations, identification of cell clusters, or sub-populations. Within scArches we allow to upload such references to a model repository via our built-in API for Zenodo (**Methods**). To enable the user to map new datasets on top of such a custom reference atlas, instead of sharing the *N* studies, we propose to share the model weights, which can now easily be downloaded from the model repository and be fine-tuned with new query (**Figure 1b**).

**Figure 1.**
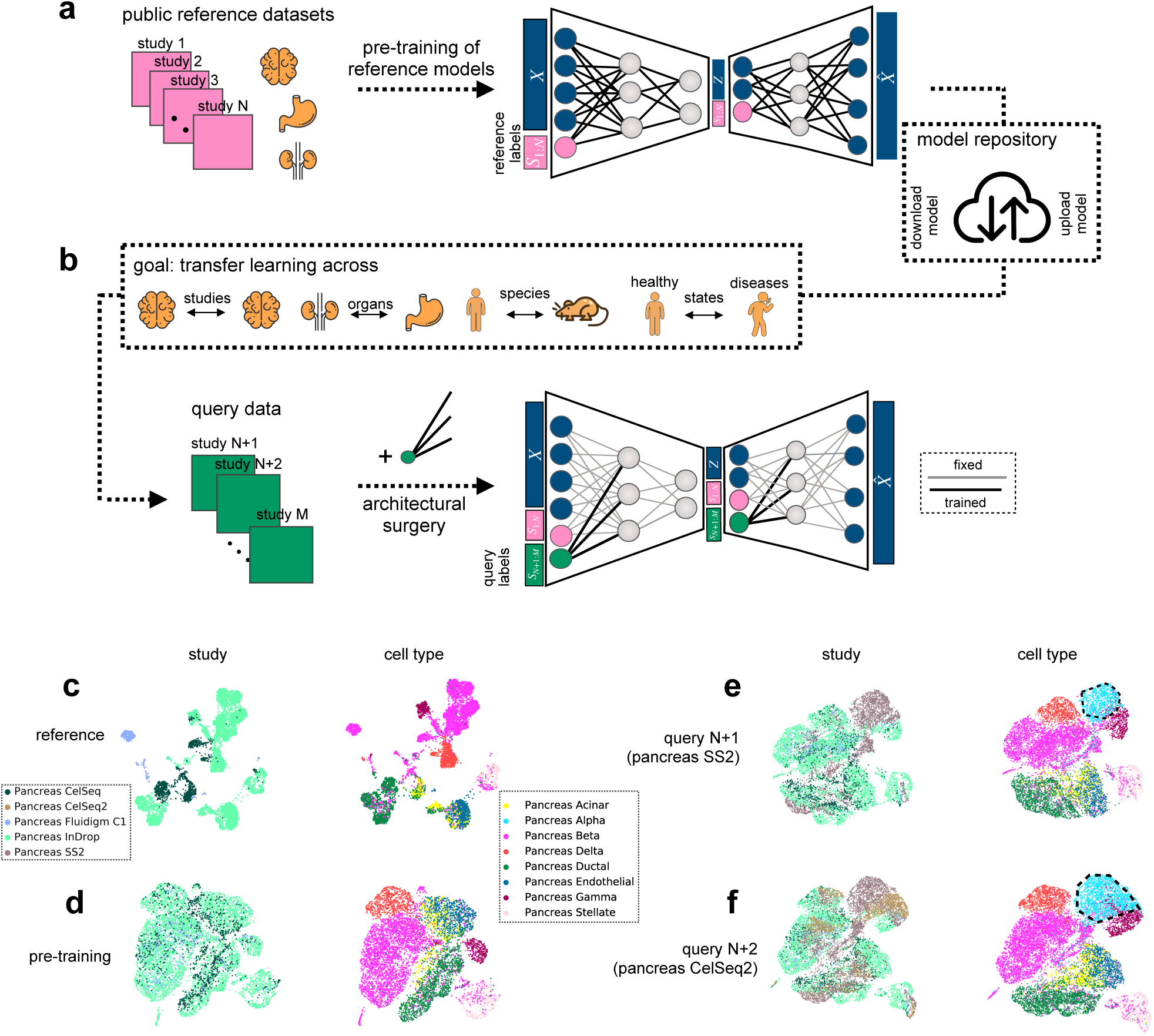
scArches enables iterative query to reference single-cell integration. **(a)** Pretraining of a latent representation using public reference datasets and corresponding reference labels. **(b)** Decentralized model building: Users download parameters for the atlas of interest, fine-tune the model and optionally upload their updated model for other users. **(c-f)** Illustration of this workflow for a human pancreas atlas. Training a reference atlas across three human pancreas datasets (CelSeq, InDrop, Fluidigm C1), UMAP embedding for the original (**c**) and the integrated reference for the pretrained models (**d**). **(e)** Querying a new SMART-Seq2 (SS2) dataset to the integrated reference. **(f)** Updating the cell atlas with a fifth dataset (CelSeq2). The black dashed circles represent cells absent in the reference data.

A potential challenge in conditional neural network models however is the fact that a study corresponds to an input neuron, which does not allow for adding new studies within the given network. To overcome this, we implement the architecture surgery approach to incorporate new study labels as new input nodes (**Methods**). As in a classical TL approach for supervised models, all weights are fully transferred to the target model without further modification. The trainable parameters of the query model however are restricted to a small subset of weights that account for the query study labels. Depending on the size of this subset (see later experiments), this restriction functions as an inductive bias to prevent the model from strongly adapting its parameters to the query studies and thus overfitting the query data. Thus, the query data updates the reference atlas. The resulting integrated data can be used for transferring cell type annotations from the reference to the query and joint analysis of reference and query data.

To illustrate the feasibility of this approach, we applied scArches with trVAE to consecutively integrate two novel studies into a pancreas reference atlas comprised of three studies, all measured with different sequencing technologies (**Figure 1c**). To additionally simulate the scenario where the query data contains a new cell type absent in the reference, we removed all the alpha cells in the training reference data. We first trained a trVAE model with scArches to integrate the training data and construct a reference atlas (**Figure 1d**). Once the reference atlas was constructed, we fine-tuned the reference model with the first query data (SS2) and subsequently updated the reference atlas with this study (**Figure 1e**). This procedure was repeated with the second query data (CelSeq2, **Figure 1f**). After each update, we observe that our model overlays all the shared cell types present in both query and reference. Additionally, reference mapping and updating yielded a separate and well-mixed cluster of alpha cells in the query datasets (black dashed circles in **Figure 1e, f**). To further assess the robustness of the approach, we held out two cell types (alpha cells and gamma cells) in the reference data while keeping both of them in the query datasets. Here, our model robustly integrates the query data while placing unseen cell types into distinct clusters (**Supplementary Figure 1**). Additional testing using simulated data showed that scArches is also robust to simultaneously updating the reference atlas with more than one query study at a time (**Supplementary Figure 2**). Overall, TL with architectural surgery enables the users to update the learnt reference models by integrating novel query data, accounting for differences in cell type composition, and missing cell types in a reference.

### Selecting fine-tuning strategies for scArches favors simple low-complexity model updates

To determine the number of weights to optimize in scArches, we evaluated the data integration performance of different fine-tuning strategies by quantifying the entropy of batch mixing (EBM) after integration (see **Methods**) [29]. While higher EBM scores indicate better mixing of cells across batches, a perfect integration score can be achieved by neglecting any biological variation and randomly mixing all the data points regardless of their cell type. To assess the preservation of small neighborhoods of cells in the original data, we added an opposing metric called k-nearest neighbors (KNN) purity (see **Methods**) [30]. A high KNN purity can be obtained by partitioning the cells into separate clusters without performing any mixing. Hence, an accurate integration algorithm would result in a high EBM score and high KNN purity (retaining the original structure of the data).

Apart from fine-tuning only the weights connecting newly added studies as proposed above, we considered two less regularized candidates for fine-tuning: (1) training input layers in both encoder and decoder while the rest of the weights are frozen, and (2) fine-tuning all weights in the model. We trained a reference model using a subset of 250, 000 cells from two mouse brain studies from Saunders *et al*. [31] and Rosenberg *et al*. [32]. Using the same reference model, we compared the integration performance of the candidate fine-tuning strategies when mapping two query datasets (Zeisel, Tabula Muris) [1, 33] onto the reference data. Applying TL with the outlined three granularity levels to the brain atlas, we find that the models with fewer parameters yield better mixing while preserving the distinctions between the different cell types (**Figure2a-c**). This can be attributed to the strong bias towards the reference model by restricting the training solely to the batch-associated weights. Specifically, the strongly regularized scArches reduces the trainable parameters by 4 − 5 orders of magnitude while also demonstrating a better mixing between the studies (**Figure 2d-e, Supplementary Figure 3**).

**Figure 2.**
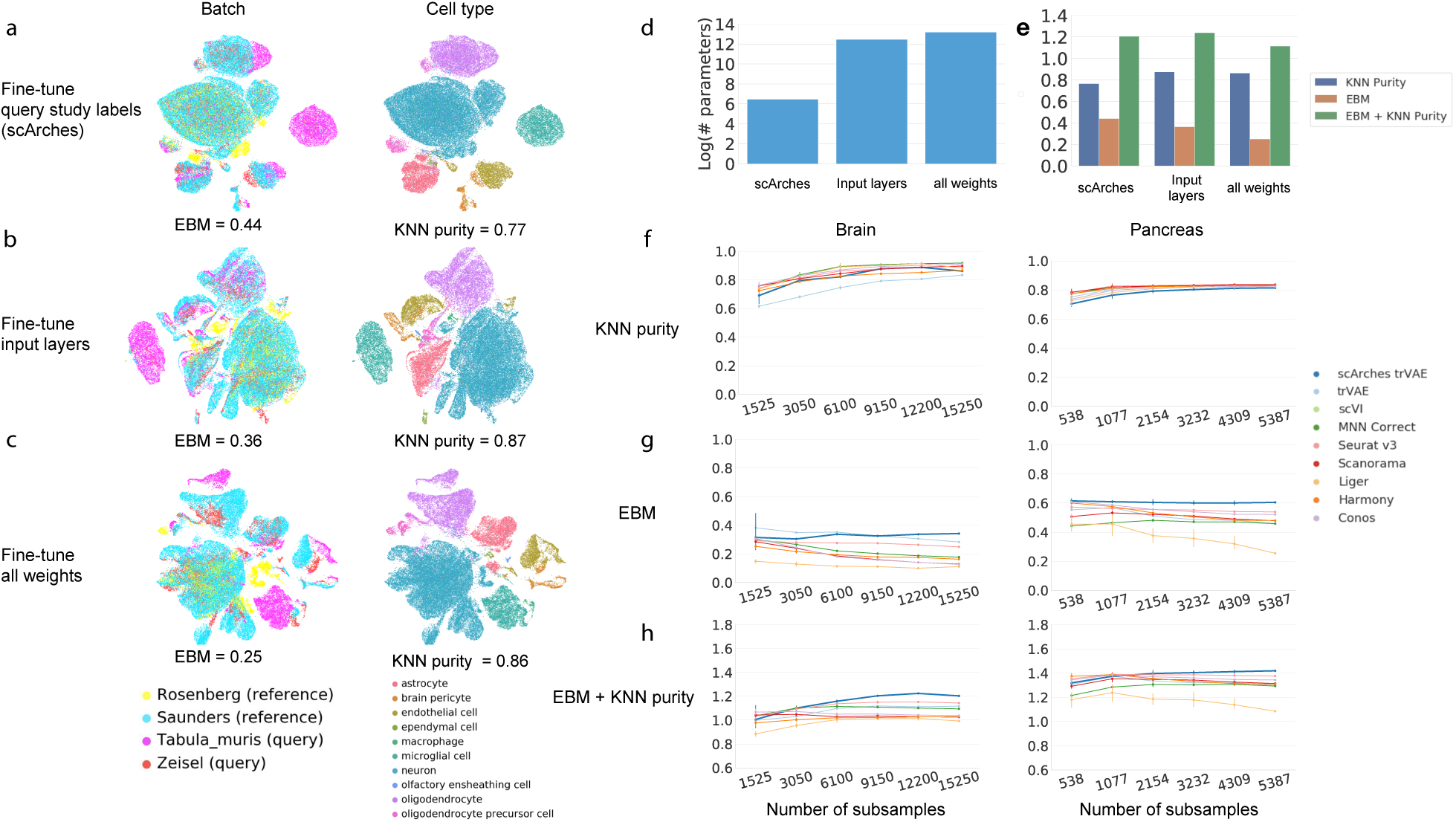
scArches enables efficient integration compared to full integration workflow with existing data integration methods. **(a-c)** Comparing different granularity levels in the proposed transfer-learning strategy by mapping two brain studies to a reference brain atlas. The reference model was trained on a subset of 250, 000 cells from two brain studies [31, 32], and then updated with Zeisel *et al*.[33] and the Tabula Muris brain subset [1]. The KNN purity (k=15) and EBM (k=15) were measured to evaluate the performances. Fine-tuning strategies vary from training a few query study labels weights **(a)** to input layers of both the encoder and decoder **(b)** or retraining full network **(c)**. The comparison of the number of the trained weights **(d)** and integration quality **(e)** on the brain atlas for these three granularity levels. **(f-h)** Evaluation of existing fully retrained integration methods versus transfer learning on the two brain (*n* = 15, 200) and pancreas (n=5, 387) datasets, respectively, for varying sample sizes. The mean and s.d. shown for n=5 random subsamples for each sample size.

### Low-complexity architectural surgery allows for efficient data integration compared to full integration methods

To demonstrate scArches’ performance in batch correction, we benchmarked our TL approach against several existing fully-trained batch-correction methods (Seurat v3 [21], Harmony [34], Liger [35], scVI [25], Scanorama [36], MNN correct [29], Conos [37], and trVAE [27]). In addition to quantifying EBM and KNN purity metrics for each method, we visualized the integrated data using UMAP (**Supplementary Figure 4-5**). We found that across 2 organs and 4 data sets, scArches+trVAE performed on par with existing methods in preserving the internal substructures of the original data, while outperforming these methods on mixing across studies. This performance gain is particularly prominent for the more challenging integration of mouse brain data sets (**Figure 2f-h**). Notably, our method substantially outperforms the baseline trVAE, which does not benefit from TL.

Additionally, we tested the effect of dataset size by assessing the quality of integration on subsamples of varying sizes. With increasing sample size, both the EBM and KNN purity increase across datasets and sample sizes when using scArches (**Figure 2f,g**). This observation only holds for scArches+trVAE and illustrates the overall performance improvement attributable to TL in large data regimes such as large cell atlases with many cells, where the robustness of mapping and cell-type identification across newly added studies is desired. Further, in the presence of low cell numbers, scArches+trVAE outperforms other existing methods in almost all cases, demonstrating the benefits of the added regularization of TL in low-data regimes in scArches. Therefore, scArches can also be used to integrate plate-based datasets that typically contain lower numbers of cells [38]. The above observations are robust to changes in kNN graph neighborhood size (**Supplementary Figure 6**). Overall, scArches improves the integration performance of DL models and outperforms other integration methods especially in large data regimes.

### Architectural surgery enables integrating cell atlases across organisms and species

Following the robust data integration performance of scArches, we investigated whether it can map queries to references across nominally stronger batch effects arising from tissues, and even species [7]. Integrating across tissues and species allows users to investigate the similarity of cellular populations across organs [2] and assess the suitability of model organisms [39, 40]. With whole-organism atlases becoming available in both mouse [1, 3] and human[4], these questions can be asked by mapping the query data on top of these references.

We considered the recently published Tabula Senis (TS) [3] as our reference, which includes 155 distinct cell types across 23 tissues and five age groups ranging from one month to thirty months from plate-based (SMART-Seq2) and droplet-based (10x Genomics) assays. As the query data, we used the cells from the 3-month time point, which is equivalent the previously published Tabula Muris (TM) dataset [1]. The query data consists of 90, 120 cells from 24 tissues including an out-of-distribution tissue, trachea, which we excluded from the reference data. trVAE with scArches accurately integrates the query and reference data across time points and sequencing technologies and creates a distinct cluster of tracheal cells (*n* = 9, 330) (**Figure 3a-c**).

Building upon the query-reference embedding, we investigated the transfer of cell-type labels from the reference dataset. We approached this classification problem by first training a simple kNN classifier on the latent space representation of the reference TS. Then each cell in the query TM was annotated using its closest neighbors in the reference dataset. Additionally, our classification pipeline provides an uncertainty score for each cell while reporting cells with more than 50 % uncertainty as unknown (see **Methods**). Our model transferred the labels from the reference atlas to the query atlas with ≈ 89% accuracy for all the tissues except tracheal cells (**Figure 3d**). Moreover, all misclassified cells and cells from the out-of-distribution tissue received high uncertainty scores (**Figure 3e-f**). Overall, the classification results across tissues indicated a robust prediction accuracy across most tissues (**Figure 3g**) while highlighting which cells were not mappable to the reference. The robust performance of a simple KNN classifier on the integrated latent space demonstrates that scArches can successfully merge large and complex query datasets into reference atlases.

**Figure 3.**
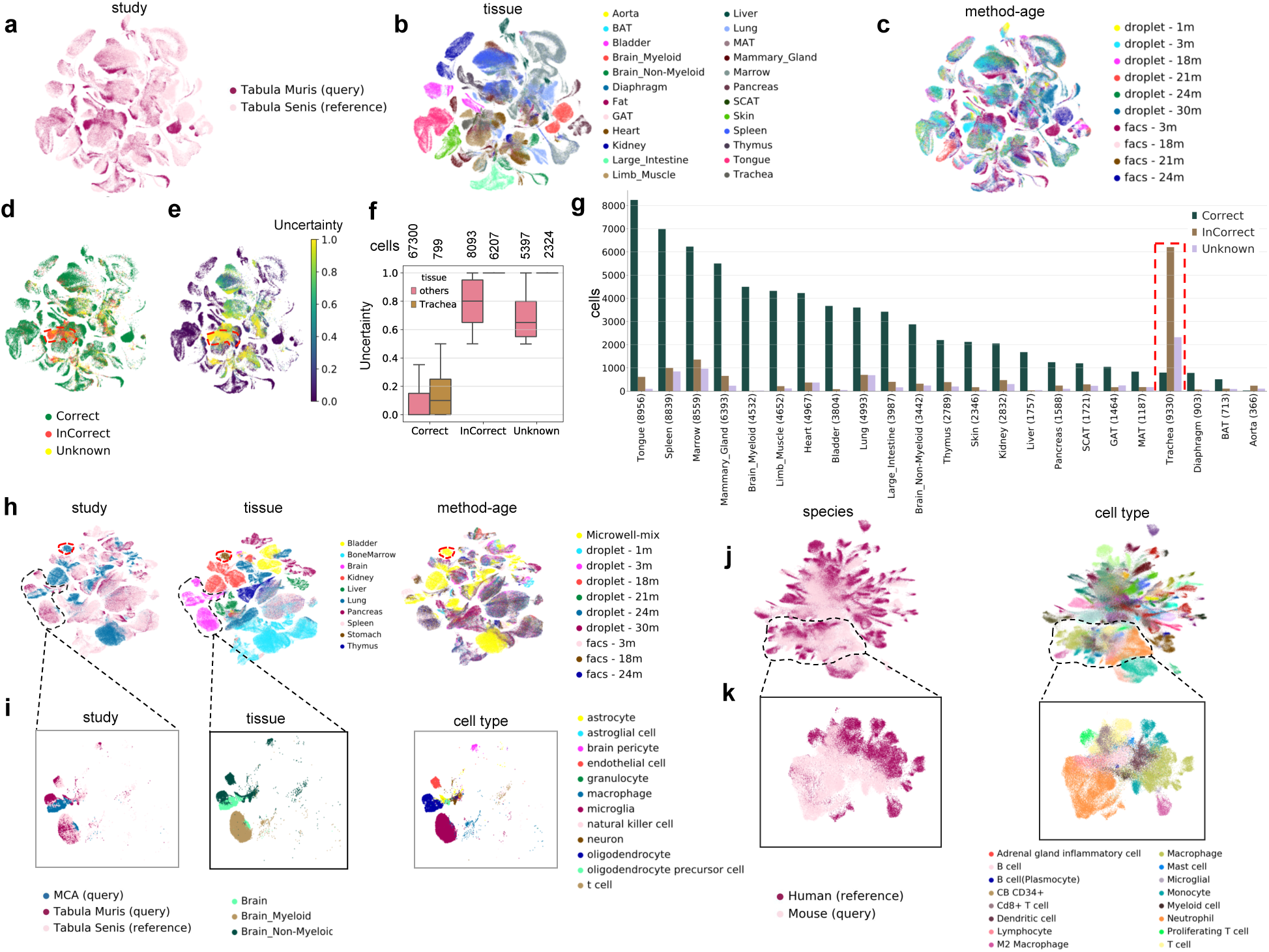
scArches successfully integrates data across whole organisms and between species. **(a-c)** Querying Tabula Muris (n=90, 120) to the larger reference atlas Tabula Senis (n=264, 287) **(a)** across different technologies, tissues **(b)**, and ages **(c)**. The tissues were correctly grouped across the two data sets **(a-b). (d)** Location of misclassified and unknown cells after transferring the labels from the reference to the query data. The highlighted tissue represents the trachea cells, which were removed from the reference data. **(e)** Reported uncertainty of the transferred labels, which was low in the correctly classified cells **(f)** and high in the incorrect and unknown ones, particularly in trachea. **(g)** The number of correct, incorrect, and unknown cells across different tissues. The red dashed line represents trachea cells only present in Tabula Muris. **(h)** Aligning query Mouse Cell Atlas (MCA, n=71, 259) and Tabula Muris (n=43, 127) mouse atlases into Tabula Senis (n=169, 425). The red dashed circle represents the stomach cells only present in MCA. **(i)** Alignment of query brain cells across both the myeloid and non-myeloid in the reference. **(j)** Querying MCA (n=122, 924) to the reference human cell atlas (n=249, 845). **(k)** The cross-species comparison between immune cells, illustrating mixedness across species.

To demonstrate the scalability and robustness of architecture surgery for whole-atlas integration, we used scArches to map a large query of both the Mouse Cell Atlas (MCA) [2] (Microwell-seq) and TM (Smart-seq2, 10X) to the TS reference atlas. As the MCA was profiled using a different technique and was sequenced at a shallow depth, integrating this dataset has been reported to be a considerably harder challenge [7].

Using atlas type and sequencing technique as batch labels, our model successfully groups similar tissues from different atlases while preserving the heterogeneity within each tissue (**Figure 3h**). Illustratively, we further examined two major brain cell clusters after integration. scArches successfully aligned the query microglia cells to the myeloid brain cell cluster in the reference while non-immune glial cells such as astrocytes and oligodendrocytes were correctly integrated to the non-myeloid cluster (**Figure 3i**). Thus, transferring weights from the TS reference model to co-embed MCA data enabled the integration of different atlases to overcome batch effects from different laboratories, technologies, and ages.

In addition to integrating studies from an organism, it is instructive to assess the similarity of cell types across species. Illustratively, we trained a reference model based on the recently published Human Cell Landscape (HCL) [4] comprised of 249, 845 cells across 63 human tissues. After architecture surgery, we aligned the MCA (n=122, 944) into the reference human cells (**Figure 3j**).

Given the abundance of species-specific cell types and species-specific functions of particular cell types, we do not expect all cell types to overlap across species. Yet, we find that specifically similar immune cell types, such as neutrophils and macrophages, were clustered together across species while species-specific cell types were placed separately (**Figure 3k**). These observations agreed with the cluster-averaging pseudo-cell analysis from the original publication [4], confirming a high similarity of the immune cells and other major cell type across both species. In all, the strong regularization of the transfer from reference via scArches allows the integration to overcome the strong species biological effect and focus on the gene expression similarity across major mammalian cell types (**Supplementary Figure 7**); we thus believe that the resulting resource can offer a basis for cross-species analysis of cell-type identity.

### Mapping cells from COVID-19 patients onto a healthy reference retains healthy cell states and captures emergence of novel, unseen, disease-associated cell types

The query-reference setup in scArches enables users to contextualize their data using a compendium of aggregated datasets. In the study of disease, the contextualization with healthy reference data is essential. To showcase how disease contextualization can be performed with scArches, we mapped a recent dataset, containing immune and epithelial cells collected via bronchoalveolar lavage from healthy controls, and patients with moderate and severe COVID-19 (*n* = 62, 469) [41] to a reference aggregated from bone marrow [42], PBMCs [43–45], and normal lung tissue [46–48] (n=154, 723; **Figure 4a-c**). To demonstrate that our approach works with other types of models, we used a NB CVAE similar to scVI here [25] (see **Methods**).

**Figure 4.**
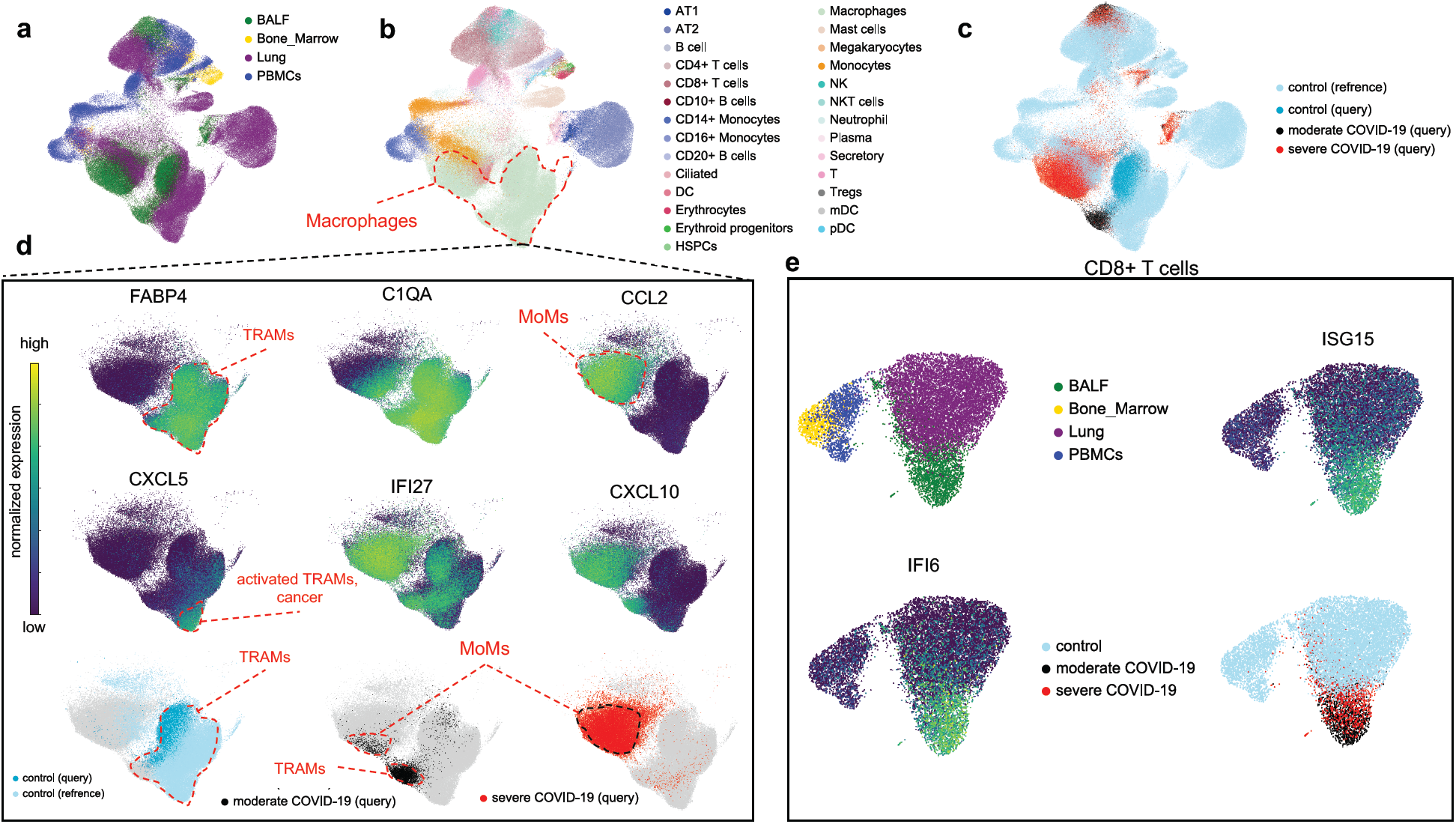
scArches resolves severity in COVID-19 query data mapped to a healthy reference and reveals emergent cell states. **(a-c)** Integration of query immune and epithelial cells from patients with COVID-19 on the top of healthy immune atlas across multiple tissues **(a)**, cell types **(b)** and cell states **(c). (d)** Comparison of various macrophage subpopulations across both healthy and COVID-19 states. Top row: Tissue-resident alveolar macrophages (TRAMs) are characterized by expression of *FAPB4*, while monocyte-derived inflammatory macrophages (MoMs) by expression of *CCL2*. Upregulation of *C1QA* illustrates maturation of MoMs as they differentiate from monocytes to macrophages. Middle row: *CXCL5, IFI27* and *CXCL10* illustrate context-dependent activation of TRAMs. Bottom row: scArches correctly maps TRAMs from query to TRAMs from reference, while preserving MoMs, unseen in the reference, as a distinct cell type. **(e)** Separations of activated query CD8+ T cells from patients with COVID-19 from the rest of CD8+ T cells in the reference.

A successful disease-to-healthy data integration should satisfy three criteria: (1) preservation of biological variation of healthy cell states; (2) integration of matching cell types between healthy reference and disease query; and (3) preservation of distinct disease variation, such as the emergence of the new cell types that are unseen during the healthy reference building. The COVID-19 query data contains the following cell types: airway epithelial cells, plasma and B cells, CD8+ T cells, neutrophils, monocytes, mast, natural killer cells, dendritic cells, and macrophages (**Figure 4b,c** and **Supplementary Figure 8**).

Within the macrophage cluster, two distinct macrophage populations dominate the structure of the embedding (**Figure 4d**): tissue-resident alveolar macrophages (TRAMs; *FABP4* + and *C1Q* +), and monocyte-derived inflammatory macrophages (MoMs; *FABP4* -, *CCL2* +). Control samples from the query dataset were obtained from healthy controls and dominated by TRAMs, which integrate well with the healthy reference data from the lung (**Figure 4c,d**). Samples from patients with moderate COVID-19 contained both TRAMs and MoMs. While TRAMs from moderate COVID-19 integrated with TRAMs from control lung tissue, they did not mix with normal TRAMs completely, as they were activated and expressed *IFI27* and *CXCL10*. Similarly, activated TRAMs (*FABP4* +, *IL1B* +, *CXCL5* +) that originate from a single subject (donor 2 in the Travaglini *et al*. dataset) also formed a distinct subcluster within TRAMs. MoMs are predominantly found in samples from patients with severe COVID-19, and to a lesser extent samples from patients with moderate COVID-19. MoMs originate from monocytes that are recruited to sites of infection and thus do not appear in healthy reference tissue. Indeed, MoMs were placed in closer proximity to monocytes than TRAMs in our embedding, reflecting their ontological relationship (**Supplementary Figure 9** for PAGA [49] proximity analysis). The monocyte to MoM differentiation is illustrated by the gradient of *C1QA* expression. Activation of CD8+ T cells is another feature of COVID-19 [40]. Subsetting CD8+ T cells from our integrated data object, we find CD8+ T cells from patients with COVID-19 separating from the lung reference cells as these are in an activated state (**Figure 4e**; interferon-stimulated activation signature *ISG15* +, *IFI6* +).

Overall, the scArches joint embedding was dominated by nuanced biological “variation, e.g., macrophage subtypes even when these subtypes were not annotated in reference datasets (e.g., activated TRAMs from patients with moderate COVID-19 or a patient with lung tumor). Although disease states were absent in the reference data, scArches successfully separated these states from the healthy reference, and even preserved disease-specific variation patterns. Hence, we found that disease-to-healthy integration with scArches met all three criteria for successful integration.

## Discussion

We introduced architectural surgery, a straight-forward approach for transfer learning, reusing neural network models by adding input nodes and weights and then fine-tuning those. Architectural surgery can extend any deep learning-based data integration method to enable decentralized reference updating, facilitate model reuse, and provide a framework for learning from reference data. With our scArches implementation of this idea for single cell transcriptomics, we reduce the complexity of the data integration process and thus allow models to scale to millions of cells and enable model sharing via a model database.

In applications, we find that joint latent representations of the query and reference data allow the discovery of rare biological states in the query dataset while facilitating the transfer of knowledge from reference to query. Specifically, we demonstrated how the integration of whole-species atlases enables the transfer of cell type annotations from a reference mouse atlas to a query atlas across 24 organs and 155 cell types. We further showed that COVID-19 query data can be mapped on top of a healthy reference while retaining variation among both disease and healthy states.

The reduction in model training complexity by architectural surgery leads to both an increase in speed as well as improved usability, since mapping a query dataset to a reference requires no further hyperparameter optimization. With scArches, one can therefore use pretrained neural network models without computational expertise or much, if any, GPU power to map e.g., disease data onto stored reference networks prepared from independent or aggregated tissue atlases. We make use of these features by providing a model database, where pretrained models are made publicly available and can be uploaded and downloaded.

An important step in using scArches is the choice of reference model. Reference building is still an unsolved problem in scRNA-seq analysis [7]. Yet, even using our imperfect reference models (see **Supplementary Note 1**), we can investigate the similarity between cell types across species or contextualize COVID-19 patient data with a healthy reference. In order to maintain optimal query integration, it is important to keep the reference model up to date. Crucially, scArches can be used with any reference built via a generative model. By promoting the sharing of reference models via our Zenodo repository we enable users to map their data onto the latest updates provided by the community.

Model sharing in combination with reference mapping via scArches allows users to create custom reference atlases by updating public ones, and paves the way for automated and standardized analyses of single cell studies. In our opinion, model sharing is a crucial next step in the dissemination of single cell genomics methods. Especially for human data, it is often difficult to share expression profiles due to data protection regulations. Indeed, a recent Human Cell Atlas study that aggregated 31 published and unpublished human lung datasets to study the distribution of SARS-CoV-2 entry gene expression across donors [40] was limited to 3 genes and few covariates due to difficulties in sharing especially unpublished data. With scArches, users can get an overview of the whole dataset to validate harmonized cell type annotation despite being limited to sharing only three genes. By sharing a pretrained neural network model that can be locally updated, international consortia can generate a joint embedding without requiring access to the full gene sets.

We envision two major directions for further application and development. Firstly, scArches can be applied to generate context-specific large-scale disease atlases. Large disease reference datasets are increasingly becoming available [50–52]. By mapping between disease references, we can assess the similarity of these diseases at the level of single cells and cell types and thus inform drug repurposing studies. The suitability of model organisms for disease research can be investigated by mapping between the relevant species-specific disease reference atlases. For example, projecting mouse single-cell tumor data on a reference human patient tumor atlas can help to identify accurate tumor models that include desired molecular and cellular properties of the microenvironment observed in patients. Secondly, the approach can be readily extended to assemble multimodal single-cell reference atlases in order to include epigenomic [53], chromosome conformation [54], proteome [55] and spatially resolved measurements.

In summary, we have shown that architectural surgery leverages transfer learning to enable model reuse. With the increasingly common availability of reference atlases for many tissues and species, we expect scArches to enable users to easily integrate new experiments on top of those references, reusing embeddings and annotations.

## Code availability

The software is available at https://github.com/theislab/scarches. The code to reproduce the results of this paper is also available at https://github.com/theislab/scarches-reproducibility.

## Data availability

All of the datasets analyzed in this manuscript are public and published in other papers. We have referenced them in the manuscript and they are downloadable at https://github.com/theislab/scArches-reproducibility.

## Author Contributions

M.L conceived the project with contributions from F.J.T. and Z.A. M.L., M.N. and M.K. implemented the models and analyzed the data. M.B. curated the mouse brain dataset. M.L and M.D.L performed analysis of the COVID-19 dataset with the help from A.V.M. F.J.T. supervised the research. All authors wrote the manuscript.

## Acknowledgments

We are grateful to all the members of the Theis lab. M.L. is grateful for valuable feedback from Alex Wolf regarding metrics selections for batch effect removal. This work was supported by BMBF grant nos. 01IS18036A and 01IS18053A, by the German Research Foundation within the Collaborative Research Center 1243, Subproject A17, by the Helmholtz Association (Incubator grant sparse2big, grant no. ZT-I-0007) and by the Chan Zuckerberg Initiative DAF (advised fund of Silicon Valley Community Foundation, no. 182835). M.L. acknowledges financial support from the Joachim Herz Stiftung.

## Methods

### scArches

#### Architecture surgery

Our method relies on a concept known as transfer learning. Transfer learning is an approach where weights from a model trained on one task are taken and used as weight initialization or fine-tuning for another task. We introduce an *Architecture Surgery*, a strategy to apply transfer learning in the context of conditional generative models and single-cell data. Our proposed method is general and can be used to perform transfer learning on both conditional variational autoencoders (CVAEs) and Conditional Generative Adversarial Nets (CGANs) [56].

Let us assume we want to train a reference CVAE model with a *d* dimensional dataset (***x*** ∈ IR^d^) from *n* different studies (***s*** ∈IR^n^). We further assume that the bottleneck ***z*** with layer size is *k* (***z*** ∈ IR^k^). Then, an input for a single cell *i* will be *x* = *x s*, where *x* and *s* are the *d*-dimensional gene expression profile and *n* dimensional one-hot encoding of study labels respectively. The · symbol denotes the row-wise concatenation operation. Therefore, the model receives a (*d* + *n*)-dimensional and (*k* + *n*)-dimensional vectors as inputs for encoder and decoder, respectively. Assuming *m* query datasets, the target model will be initialized with all the parameters from the reference model. To incorporate *m* new study labels, we add *m* new dimensions to *s* in both the encoder and decoder networks. We refer to these new added study labels as *s′*. Next, the *m* new randomly initialized weight vectors are also added to the first layer of the encoder and decoder. Finally, we fine-tune the new model by only training the weights connected to the last *n* +*m* dimensions of *x* that correspond to the condition labels. Let us assume that *p* and *q* are the number of neurons in the first layer of the encoder and decoder, then during the fine-tuning only (*n* + *m*) *×* (*p* + *q*) parameters will be trained. Let us parameterize the first layer of the encoder and decoder part of the scArches as *f*_1_ and *g*_1_, respectively. Let us further assume that ReLU activations are used in the layers. So the equations for *f*_1_ and *g*_1_ are:

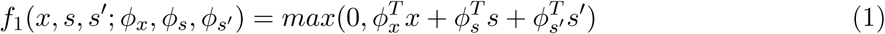

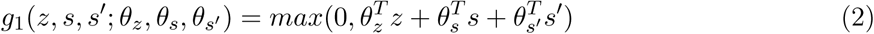

Therefore, the gradients of f and g with respect to *ϕ*_*s′*_ and *θ*_*s′*_ are:

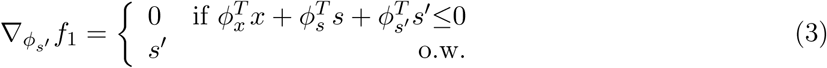

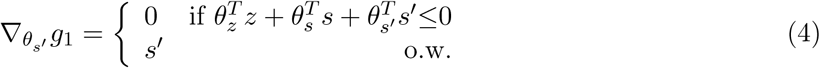

Finally, since all the other weights except *ϕ*_*s′*_ and *θ*_*s′*_ are frozen, we only compute the gradient of the scArches’s cost function with respect to *ϕ*_*s′*_ and *θ*_*s′*_ :

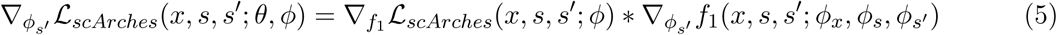

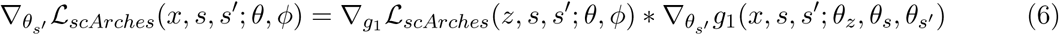

### Transfer Variational autoencoders

Variational autoencoders (VAE) were shown to learn the underlying complex structure of data. The trVAE builds upon VAE [57] framework with a motivation for providing a solution for the variational inference using neural networks to maximize the following equation:”

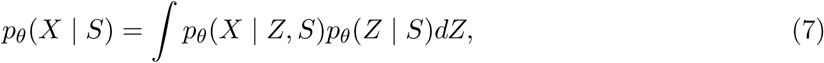

where *X* is a random variable representing the model’s input, *S* a random variable indicating various conditions, *θ* the neural network parameters, and *p*_*θ*_(*X* | *Z, S*) the output distribution that we sample *Z* to reconstruct *X*. In the following we exploit notations from [28] and a tutorial from [58]. We approximate the posterior distribution *p*_*θ*_(*Z* | *X, S*) using the variational distribution *q*_*ϕ*_(*Z* | *X, S*) that is approximated by a deep neural network parameterized with *ϕ*:

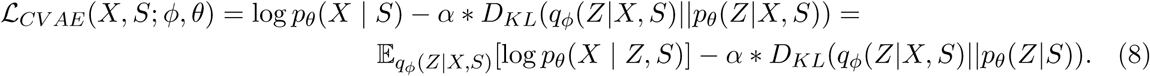

Where *θ* = {*θ′, θ*_*z*_, *θ*_*s*_} and *ϕ* = {*ϕ′, ϕ*_*x*_, *ϕ*_*s*_} are parameters of decoder and encoder, respectively. On the left-hand side, we have the log-likelihood of the data and an error term that depends on the capacity of the model.The right hand side of 8 is also known as the evidence lower bound (ELBO). Conditional variational autoencoder (CVAE) [59] is an extension of VAE framework in which *S* ≠ ø. Following the proposed method by Lotfollahi *et al*.[27], we use the representation of the first layer in the decoder, which is regularized by maximum mean discrepancy [60]. For the implementation, we use multi-scale RBF kernels defined as:

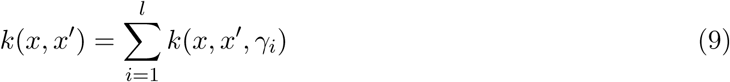

where 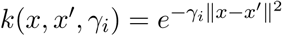 and *γi* is a hyper-parameter.

We will parameterize the encoder and decoder part of scArches as *f*_*ϕ*_ and *g*_*θ*_, respectively. So the networks *f*_*ϕ*_ and *g*_*θ*_ will accept inputs ***x, s*** and ***z, s***, respectively. Let us distinguish the first 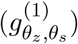 and the remaining layers 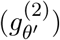 of the decoder network 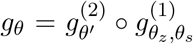. Therefore, we can define the following MMD cost function:

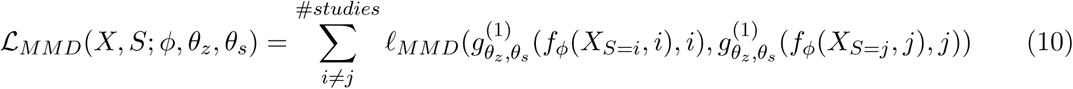

where:

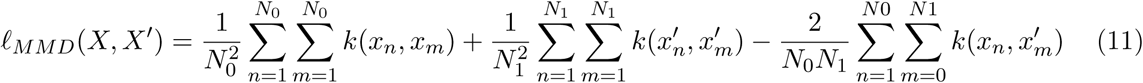

We used the notation *X*_*S*=*i*_ for the samples drawn from i-th study distribution in the training data. Finally, the trVAE’s cost function is:

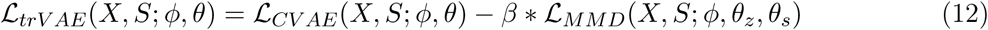

The gradients of the trVAE’s cost function with respect to *ϕ*_*s*_ and *θ*_*s*_ are:

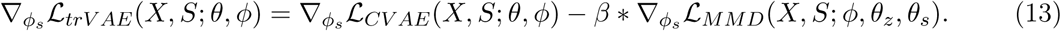

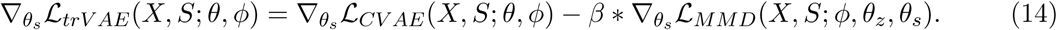

So ℒ_*trVAE*_ can be optimized using stochastic gradient ascent with respect to *ϕ*_*s*_ and *θ*_*s*_ since all the other parameters are frozen.

### Model sharing

We currently support an API to upload and download model weights and data (if available) using Zenodo. Zenodo is a general-purpose open-access repository developed to enable researches to share datasets and software. We have provided step-by-step guides to whole pipeline from training and uploading models to download, update the model and further share them. These tutorials can be found in scArches github repository (https://github.com/theislab/scarches).

### Evaluation metrics

The evaluation metrics and their definition in the current paper were taken from the work by Xu *et al*. [30]. In all of the experiments for the batch-removal evaluation, *k* was set to 15 for KNN-graph calculation.

### K-nearest neighbor purity

This metric works by constructing two similarity matrices for the first batch using the euclidean distance on the latent representation: one matrix only includes the cells from the first batch and the other one from cells in both batches. Next, we report the average ratio of the intersection of knn-graph from each similarity matrix over their union. Similarly, we calculate the same value for the next batch and finally report mean of these two values.

### Entropy of batch mixing

This metric works by constructing a fix similarity matrix for the cells. The entropy of mixing in a region of cells with c batches is defined as:

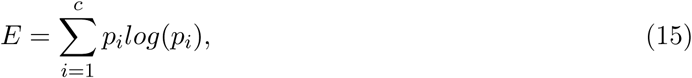

Where *p*_*i*_ is defined below as:

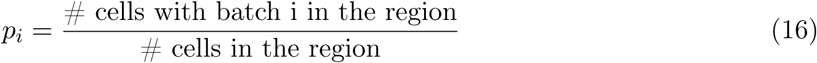

Next, we define *U*, a uniform random variable on the cell population. Let, *B*_*U*_ be frequencies of 15 nearest neighbors for the cell U in batch x. We report entropy of this variable and then average across T = 100 measurements of U.

## Datasets

### Brain data

The mouse brain dataset is a collection of four publicly available scRNA-seq mouse brain studies[1, 31–33], where additional information on cerebral regions was provided. We obtained the raw count matrix from Rosenberg et al.[32] from GEO accession ID GSE110823, the annotated count matrix from Zeisel et al.[33] from http://mousebrain.org (file name L5_all.loom, downloaded on 9/9/2019), and the count matrices per cell type from Saunders et al.[31] from http://dropviz.org (DGE by Region section, downloaded on 30/8/2019). FACS-sorted mouse brain tissue data (myeloid and non-myeloid cells, including annotation file annotations_FACS.csv) from Tabula Muris were obtained from https://figshare.com (retrieved 14/2/2019).

We harmonized cluster labels via fuzzy string matching and attempted to preserve the original annotation as far as possible. Specifically, we annotated 10 major cell types (neuron, astrocyte, oligodendrocyte, oligodendrocyte precursor cell, endothelial cell, brain pericyte, ependymal cell, olfactory ensheathing cell, macrophage and microglia). In the case of Saunders et al.[31], we facilitated the additional annotation data table for 585 reported cell types (annotation.BrainCellAtlas_Saunders_version_2018.04.01.txt retrieved from http://dropviz.org on 30/8/2019). Among those, some cell types were annotated as ‘endothelial tip’, ‘endothelial stalk’ and ‘mural’. We examined the subset of the Saunders et al. dataset as follows: We used Louvain clustering (default resolution parameter 1.0) to clusters, followed by gene expression profiling via rank_genes_groups function in scanpy. Using marker gene expression, we assigned microglia (C1qa), oligodendrocytes (Plp1), astrocytes (Gfap, Clu) and endothelial cells (Flt1) to the subset.

Finally, we applied scran normalization[61] and log(counts + 1) to transform the count matrices. In total, the dataset consists of 978, 734 cells.

### Pancreas

Five publicly available pancreatic islet datasets [62–66], with a total number of 15,681 cells in raw count matrix format were obtained from the Scanorama [36] dataset, which has already assigned its cell types using batch corrected gene expression by Scanorama [36]. The Scanorama dataset were downloaded from here. In the preprocessing step, the raw count datasets were normalized and log transformed by scanpy preprocessing methods. The preprocessed data were used directly for the pipeline of scArches.

### HCL

Human Cell Landscape (HCL) dataset was obtained from here. Raw count matrix data for all tissues were aggregated. A total number of 277,909 cells were selected and processed using scanpy python package. The data was normalized using size factor normalization such that every cell has 10,000 counts and then log transformed. Finally, 5,000 highly variable genes were selected as per their average expression and dispersion. We used the processed data directly for training the scArches at the pretraining phase.

### MCA

Mouse cell atlas (MCA) dataset was obtained from here. Raw count matrix data for all the tissues were aggregated together. A total number of 150,126 cells were selected and processed using the scanpy python package. Homologous genes were selected using BioMart 100 before merging with HCL data. The data were normalized together with HCL as explained before.

### COVID-19

COVID-19 dataset along its meta-deta were downloaded from here and here. The data set that used in this paper includes *n* = 62, 469 cells. Lung [46–48], PBMCs [43–45] and bone marrow [42] were later merged with COVID-19 samples. The data was normalized using scanpy and 2000 HVGs were selected for training the model.

### Tabula Muris Senis

Tabula Muris Senis dataset with GSE132042 as the GEO accession number is publicly available from here. The dataset contains 356,213 cells with cell type, tissue, and method annotation. We normalized the data using size factor normalization with 10,000 counts for each cell. Then, we log+1 transformed the dataset and selected 5000 highly variable genes as per their average expression and dispersion. All the preprocessing steps were done using the scanpy [67] python package. In this study, we used a combination of sequencing technology and time point as batch covariates.

### Benchmarks

We ran PCA with 10 principal components on the final results of Seurat, Scanorama, Liger, and mnnCorrect compare them to models which had latent representation.

- Harmony: We used the HarmonyMatrix function from the harmony package. We provided the function with a PCA matrix with 10 principal components on the gene expression matrix.
- Scanorama: We used the correct_scanpy function from the scanorama package with default parameters.
- Seurat: We applied Seurat as the walkthrough with default parameters.
- Liger: We used the Liger method as the walkthrough. We used k=20, lamda=5, and resolution=0.4 with other default parameters. We only scaled the data since we had already preprocessed the data.
- Conos: We followed the Conos tutorial from here. Unlike the tutorial we used our own preprocessed data for better comparisons. We used the PCA space with parameters k=30, k.self=5, ncomps=30, matching.method=‘mNN’, and metric=‘angular’ to build the graph. We set the resolution to 1 to find communities. Finally we saved the corrected psudo-PCA space with 10 components.
- mnnCorrect: We used the mnnCorrect function from the scran package with default parameters.
- scVI: We trained the scVI with symmetric architecture, having two hidden layers with 128 neurons each. This architecture yielded a better result than one hidden layer default architecture. Other parameters were set as default.

### Cell-type annotation

To classify the labels for the query dataset, We trained a weighted KNN classifier on the latent space representation of reference dataset. For each query cell *c*, we extracted its *k* nearest neighbors (*N*_*c*_). We computed the standard deviation of the nearest distances:

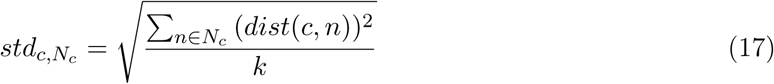

Where *dist*(*c, n*) is the euclidean distance of the query cell *c* and its neighbors *n* in the latent space. Then, we applied the Gaussian kernel to distances using:

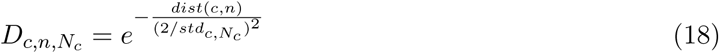

Next, we computed the probability of assigning each label *y* to the query cell *c* by normalizing across all the adjusted distances using:

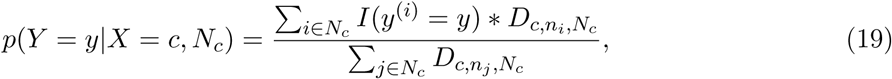

where *y*^(*i*)^ is the label of i-th nearest neighbor. Finally, we calculated the uncertainty *u* for each cell c in the query dataset using its set of closest neighbors in the reference dataset (*N*_*c*_). We defined the uncertainty *u*_*c,y*_,*N*_*c*_ for a query cell c with label y, and *N*_*c*_ as its set of nearest neighbors as:

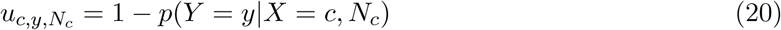

We reported cells with more than 50% uncertainty as unknown in order to detect out-of-distribution cells with new labels, which do not exist in the training data. Therefore, we labeled each cell c in the query dataset as follows:

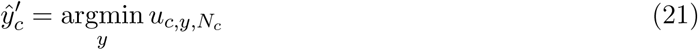

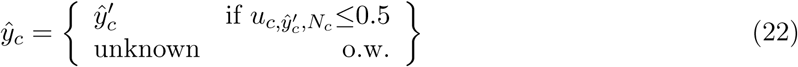

## Hyperparameters

**Table 1.** scArches detailed hyper-parameters

| model | loss | architecture | Z | $\alpha$ | $\beta$ | $\eta$ | dropout | batchnorm | clip grad |
| --- | --- | --- | --- | --- | --- | --- | --- | --- | --- |
| scArches trVAE | MSE | [128, 20] | 10 | 0.001 | 20 | 1000 | 0.05 | × | 5 |
| scArches CVAE | NB | [128, 128] | 10 | 0.0002 | 0 | 1 | 0.1 | ✓ | 3 |
| scArches trVAE | SSE | [128, 64, 20] | 15 | 0.0001 | 50 | 1 | 0.1 | × | 10 |

## 1 Supplementary Note 1

### The importance of choosing the right reference

To contextualize query data from COVID-19 patients, we generated a healthy reference dataset from lung, PBMCs, and bone marrow immune and epithelial cells. These tissues were chosen as they contain similar cell types and thus may help to contextualize the cells that we find in the COVID-19 patient data. Given that the focus of this paper is not optimal reference building, we merged these datasets using a previously well-evaluated model class (NB CVAE from scVI [28]) [7]. However, evaluating the results of mapping the COVID-19 query data into the reference model is complicated by a lack of harmonized cell type labels (e.g., some datasets have cells labeled T cells, while others separate into T cell subtypes) and the quality of the reference. These aspects are particularly complicated for the data from the Human Cell Landscape [4], which have coarse labels and do not integrate well with the rest of the data (**Supplementary Figures 8**,**10**). For example, cells with the label “T” from Liao et al seem to be CD4+ T cells (see violin plot in **Supplementary Figure 10**), but have the same label as T cells without CD4 or CD8 expression from the HCL dataset (**Supplementary Figure 10**). Indeed, our integration shows that query “T” cells map to CD4+ T cells, while the “T” cells form the HCL separate by tissue and fail to integrate into the reference. Without the effect of the HCL dataset, T cells clearly separate into CD4+ and CD8+ expressing cells across tissues and unharmonized labels. Overall, our simple reference contained sufficient relevant cells to cache the effect of the poor HCL reference. However, this may not always be the case for references used to contextualize query data. Thus, choosing the appropriate reference is of critical importance for reference mapping using scArches.

---

**Supplementary Figure 1.**
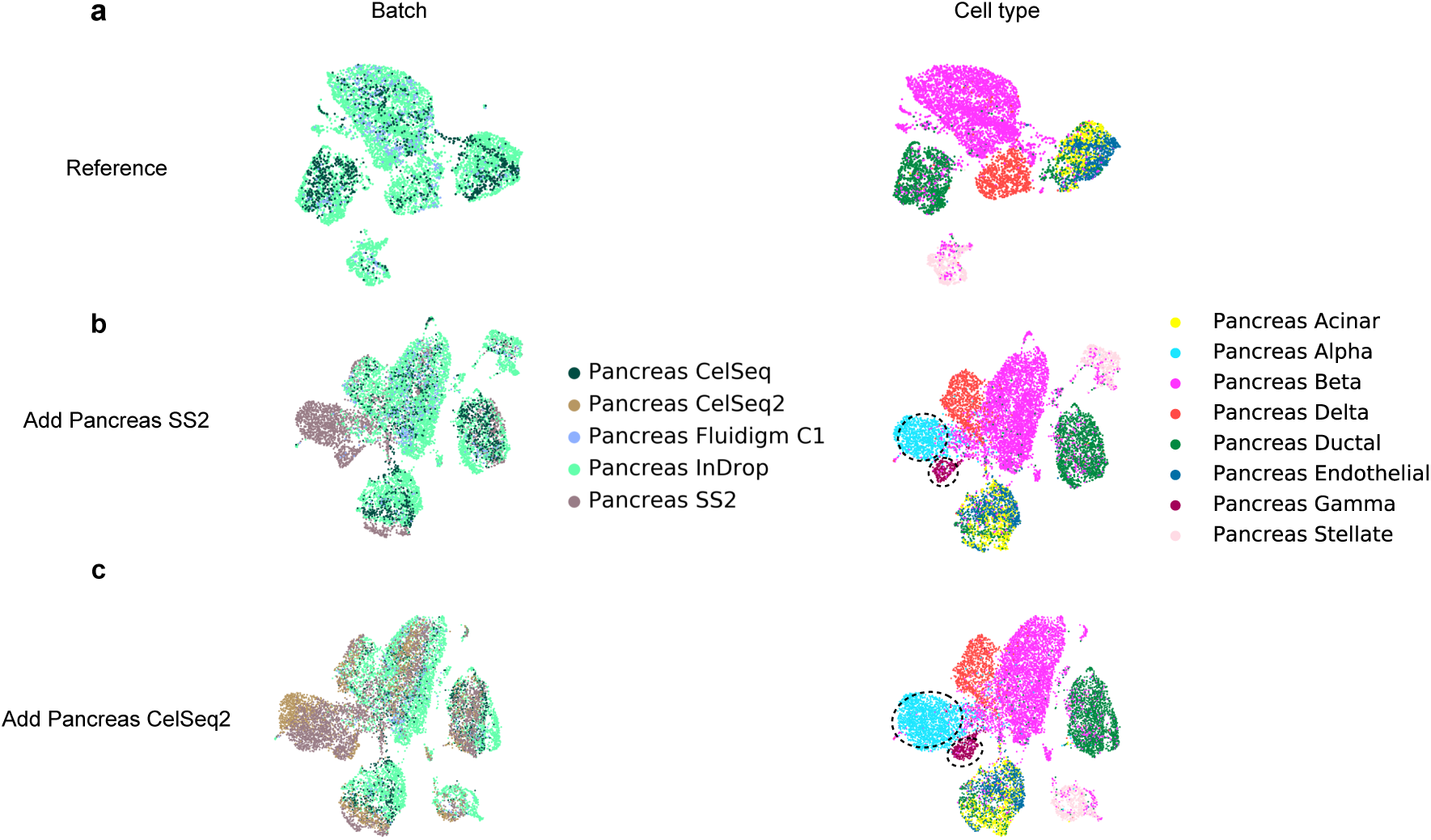
Robustness assessment for out-of-distribution cell types. **(a)** UMAP representation of the pretrained model with reference pancreas datasets (CelSeq, inDrop, Fluidgram C1) while Alpha and Gamma cells are absent in the data. **(b-c)** Iterative integration of two query data (SS2, CelSeq2) containing alpha and gamma cells by transferring weights from the reference model containing no alpha cells.

**Supplementary Figure 2.**
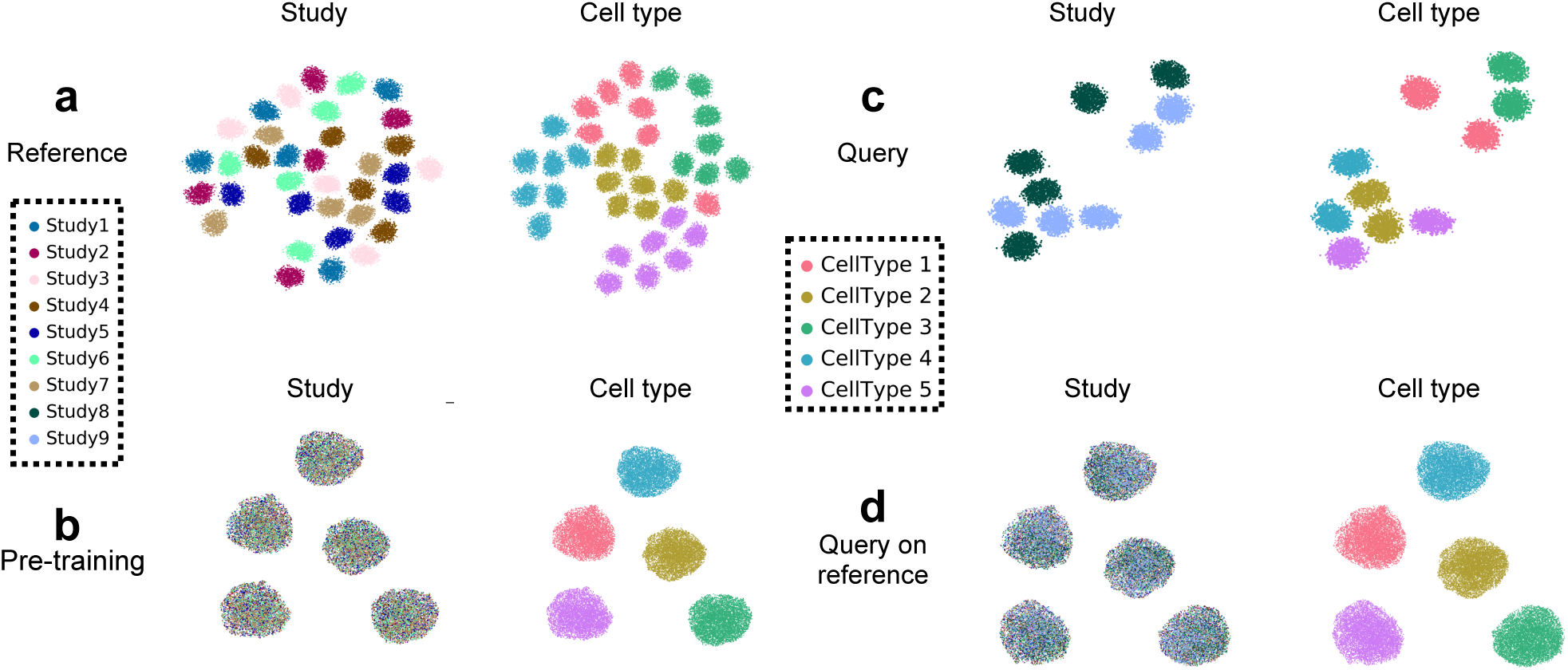
The scArches applied on simulated data. **(a)** UMAP representation of the simulated reference data with seven batches and five different cell types. **(b)** Pretraining of the model on the reference data. **(c)** The simulated query data with two batches. **(d)** Mapping the query to the reference.

**Supplementary Figure 3.**
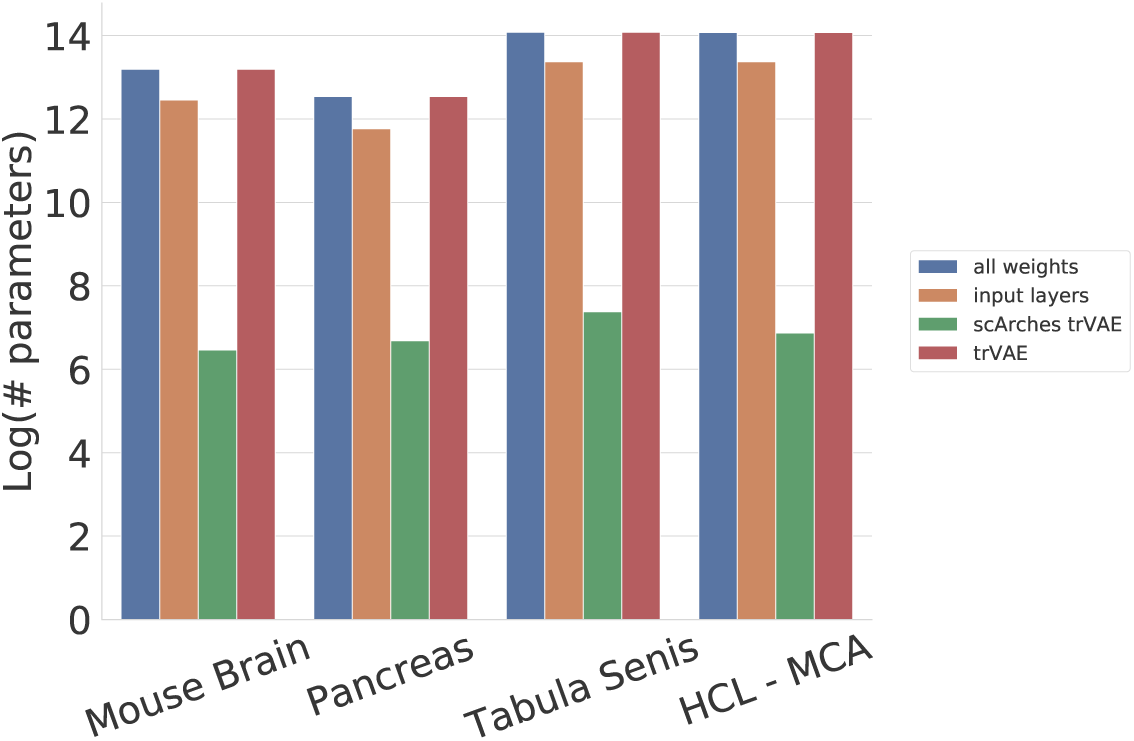
Trainable parameters comparison. The number of trainable parameters for the transfer learning models and trVAE as the baseline non-transfer learning alternatives. The vertical axis shows the number of trainable parameters in log-scale while the horizontal axis depicts different data sets.

**Supplementary Figure 4.**
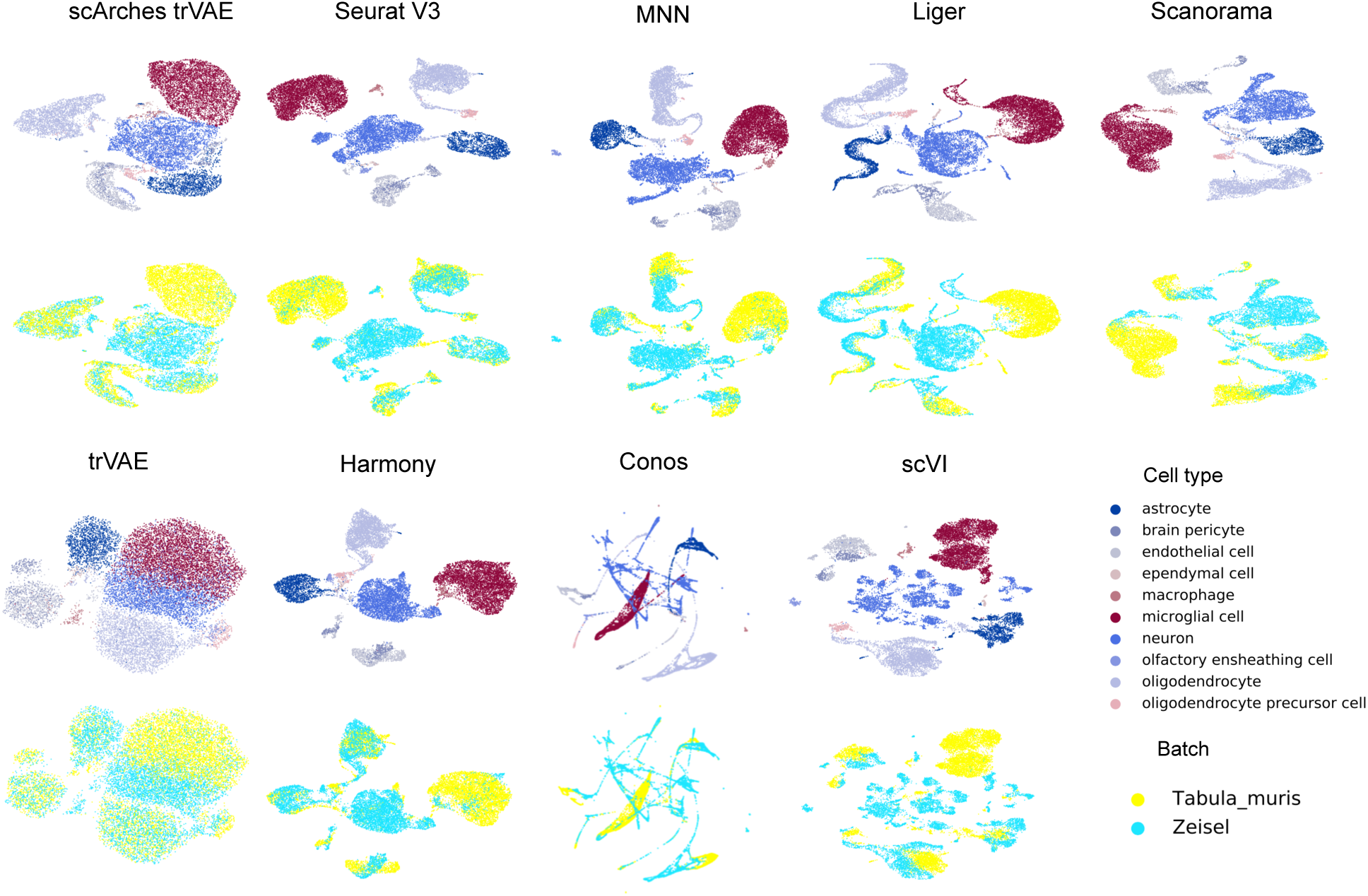
Visualisation of integrated data to compare batch-correction performance for mouse brain. Force Atlas 2 (Conos) and UMAP (all other methods) representations for brain cells after integration.

**Supplementary Figure 5.**
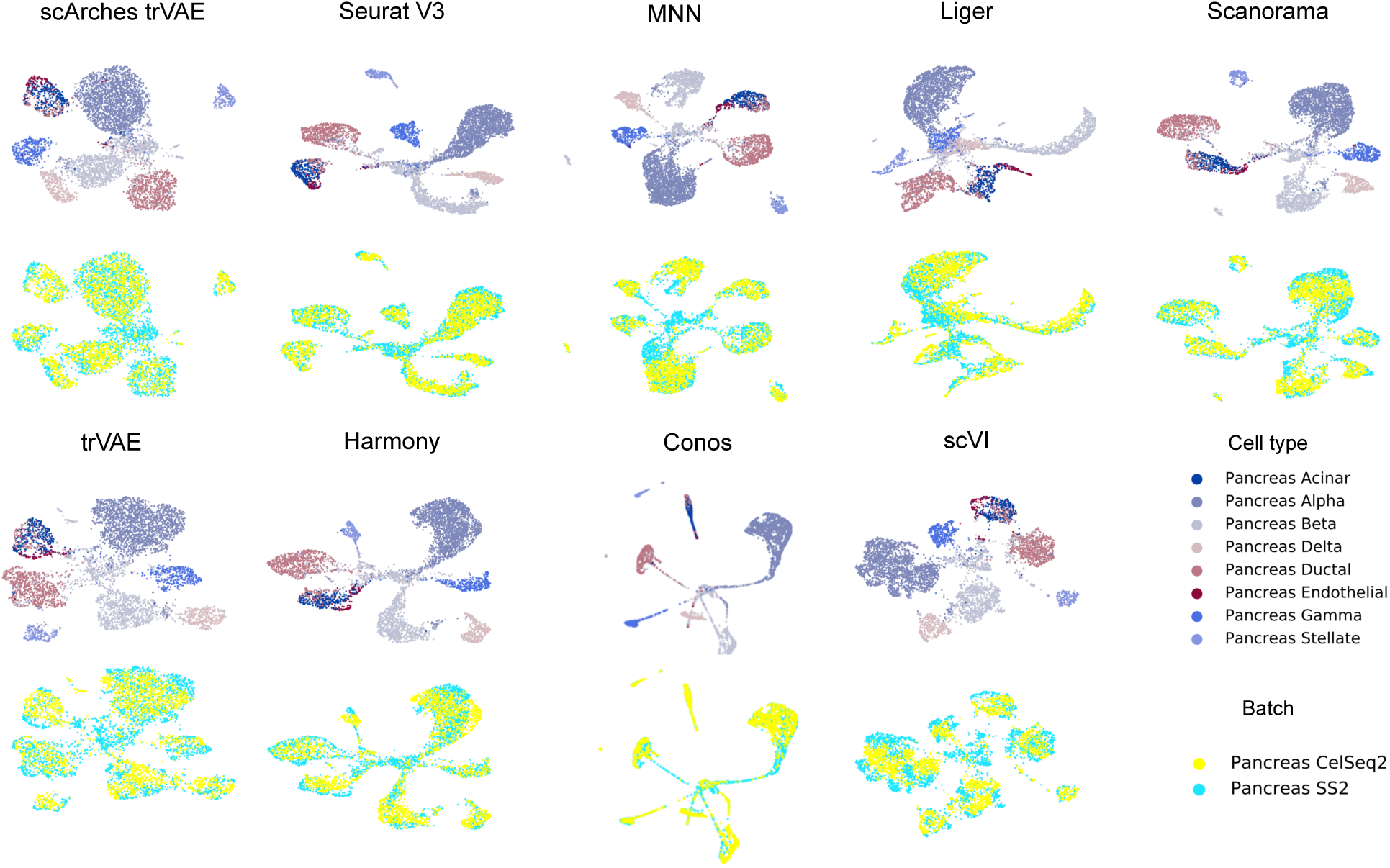
Visualisation of integrated data to compare batch-correction performance for human pancreas. Force Atlas 2 (Conos) and UMAP (all other methods) representations for pancreas cells after integration.

**Supplementary Figure 6.**
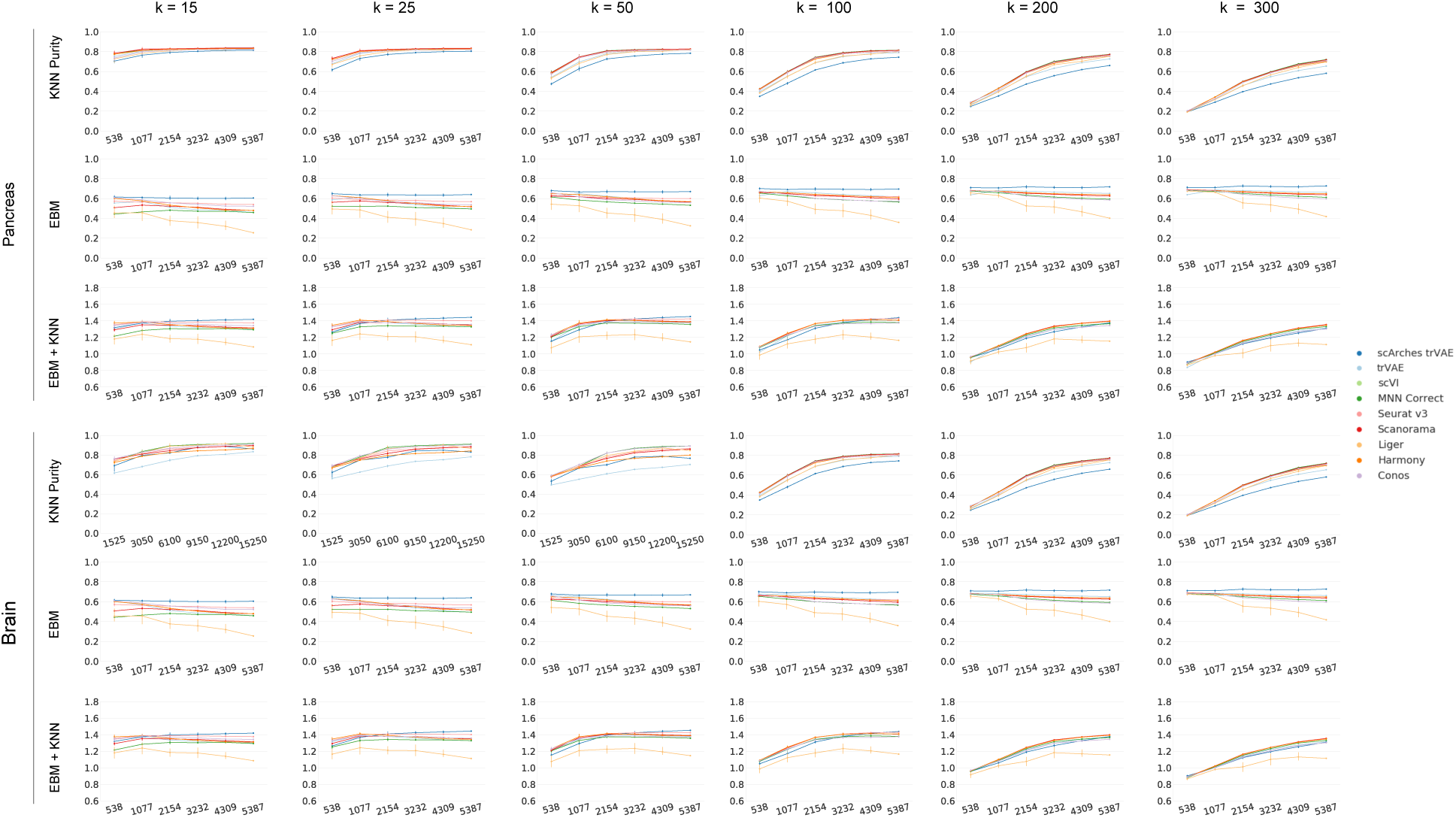
Metric comparison across different size of neighborhood (k) from high to low resolution.

**Supplementary Figure 7.**
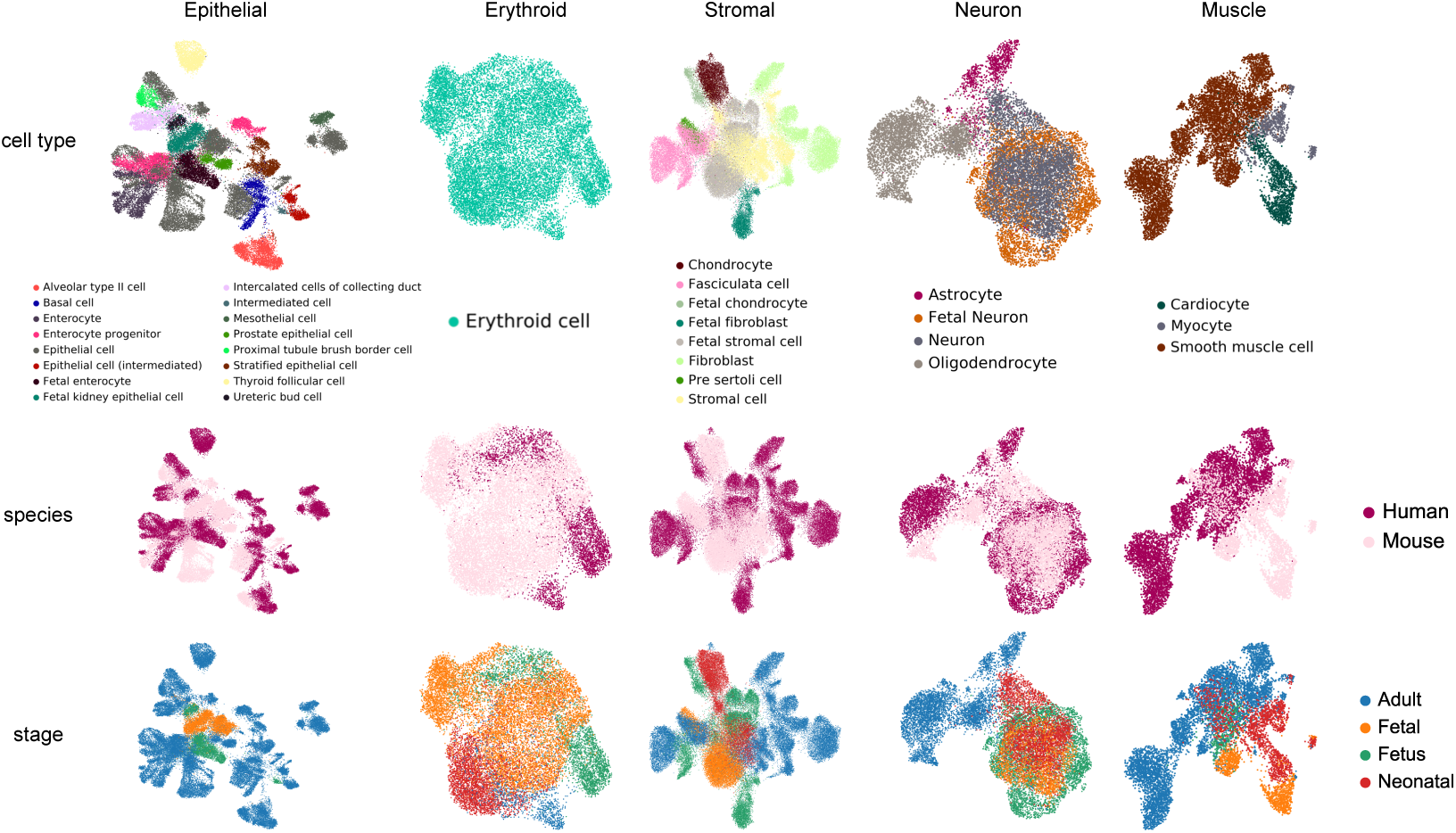
UMAP plots of major mammalian cell types across mouse and human.

**Supplementary Figure 8.**
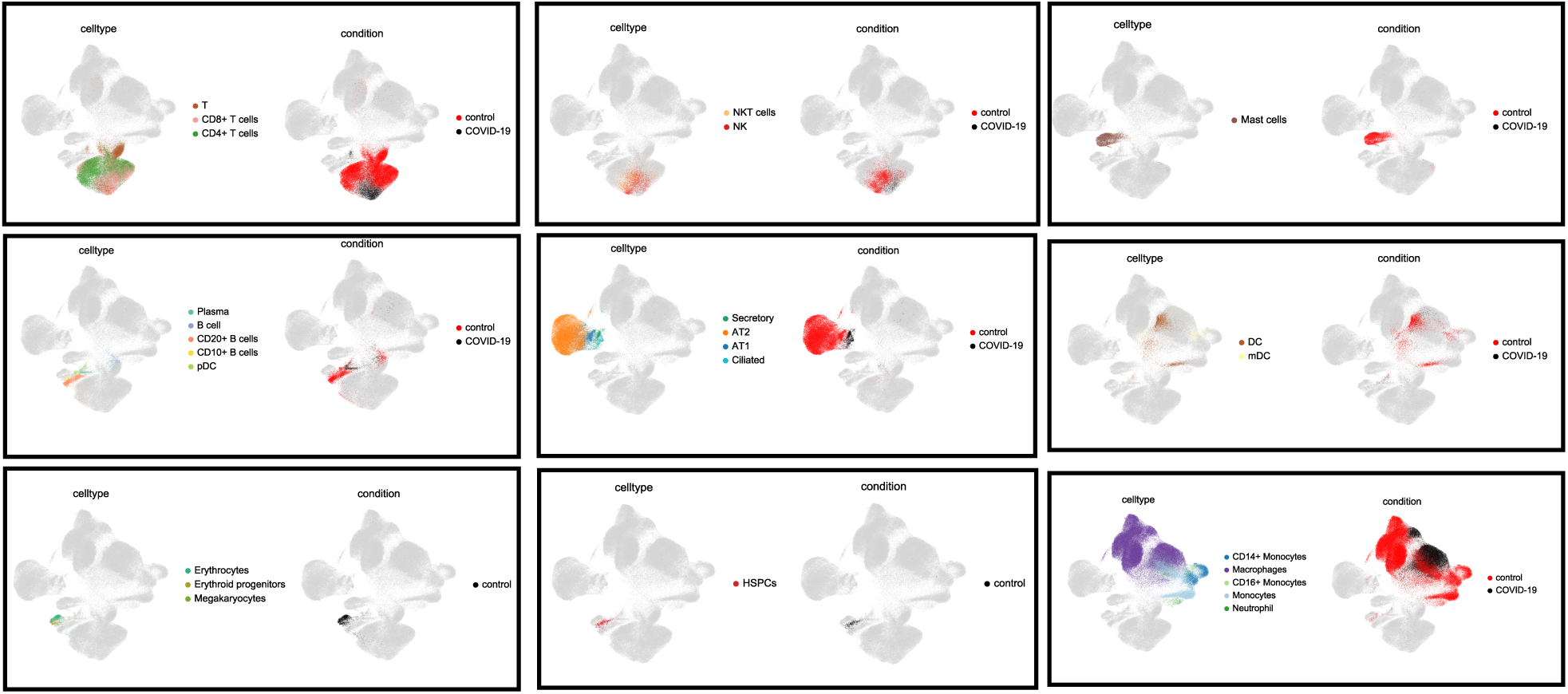
UMAP plots for all cell types present in integrated query and reference data in COVID-19.

**Supplementary Figure 9.**
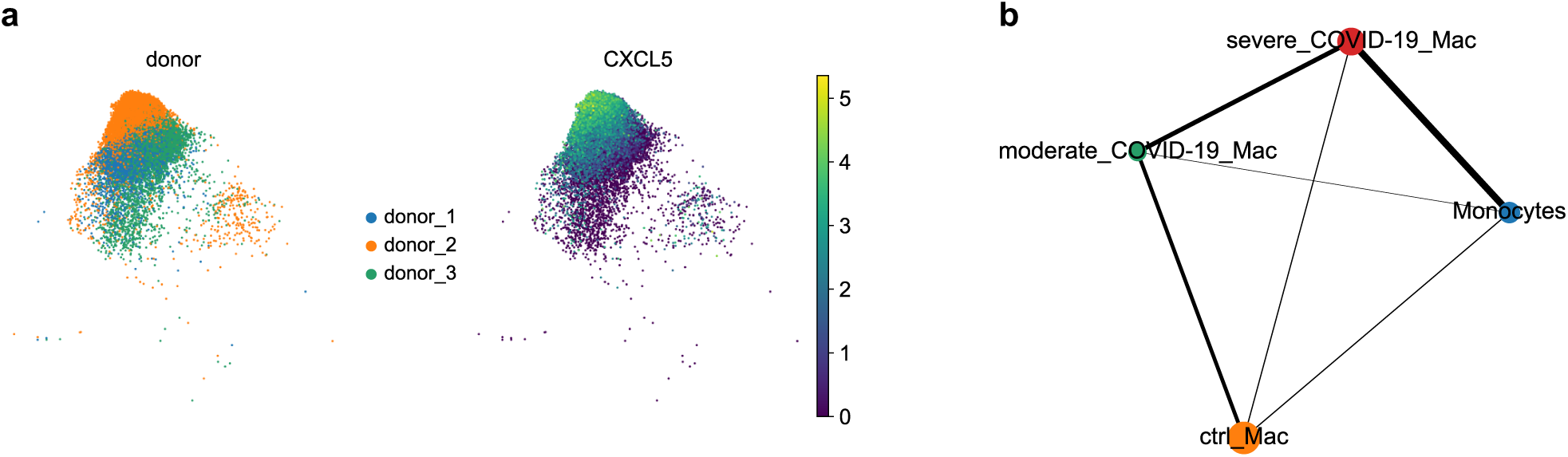
Relationship of monocyte-derived macrophages (MoMs) and monocytes in COVID-19 patients compared to a healthy reference. **(a)** cancer cells enriched with *CXLC5* from donor 2 in the Travaglini et al. dataset. **(b)** PAGA graph for monocyte and macrophage populations. Each node represents a cell state whose edge weights (represented as line thickness) quantify the connectivity between groups.

**Supplementary Figure 10.**
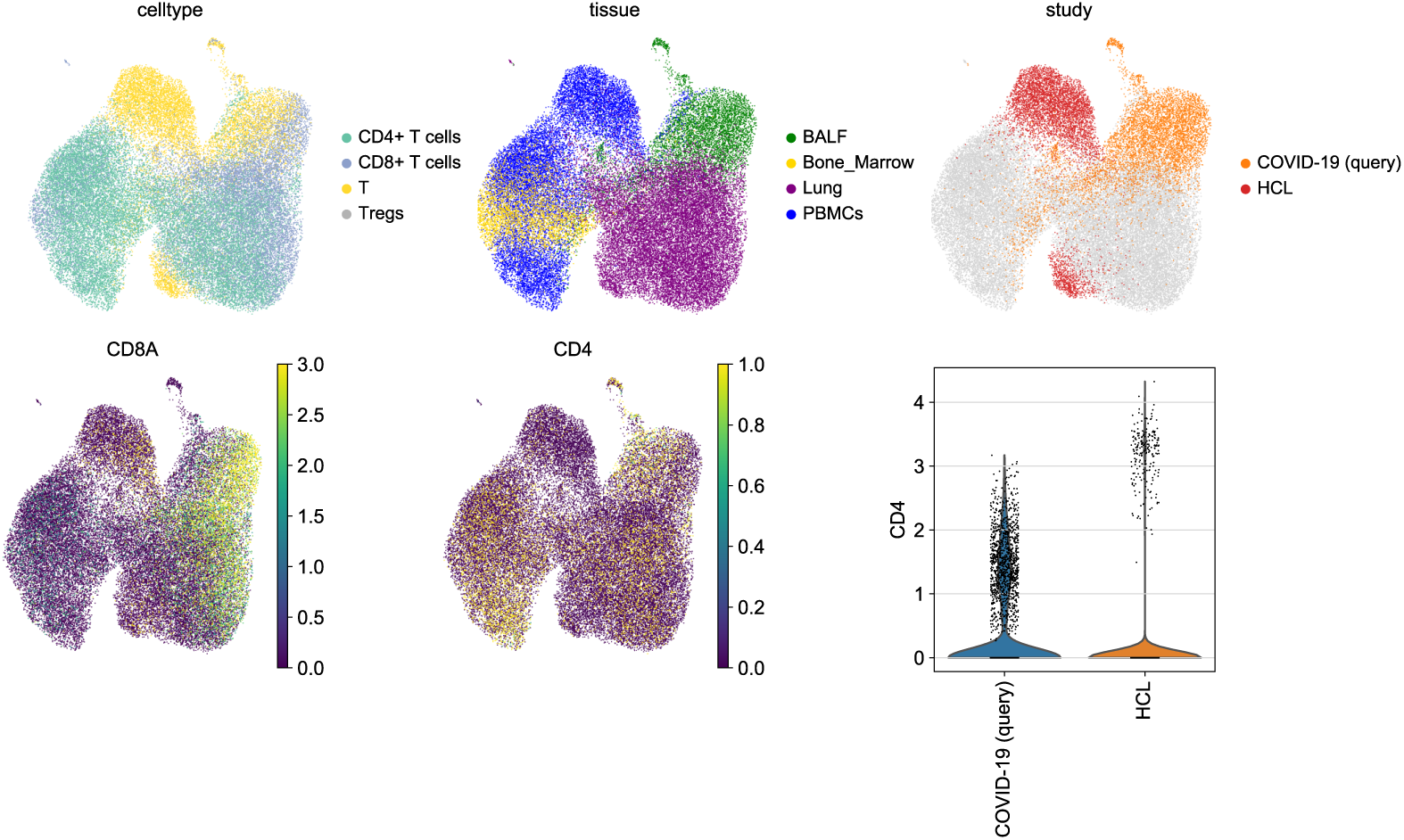
Overlap of T cell labels of all studies. Resolving T cell labels in to CD8+ T and CD4+ T cells in HCL [4] and COVID-19 datasets.

